# SARS-CoV-2 Nucleocapsid protein attenuates stress granule formation and alters gene expression via direct interaction with host mRNAs

**DOI:** 10.1101/2020.10.23.342113

**Authors:** Syed Nabeel-Shah, Hyunmin Lee, Nujhat Ahmed, Edyta Marcon, Shaghayegh Farhangmehr, Shuye Pu, Giovanni L. Burke, Kanwal Ashraf, Hong Wei, Guoqing Zhong, Hua Tang, Jianyi Yang, Benjamin J. Blencowe, Zhaolei Zhang, Jack F. Greenblatt

## Abstract

The COVID-19 pandemic has caused over one million deaths thus far. There is an urgent need for the development of specific viral therapeutics and a vaccine. SARS-CoV-2 nucleocapsid (N) protein is highly expressed upon infection and is essential for viral replication, making it a promising target for both antiviral drug and vaccine development. Here, starting from a functional proteomics workflow, we initially catalogued the protein-protein interactions of 21 SARS-CoV-2 proteins in HEK293 cells, finding that the stress granule resident proteins G3BP1 and G3BP2 copurify with N with high specificity. We demonstrate that N protein expression in human cells sequesters G3BP1 and G3BP2 through its physical interaction with these proteins, attenuating stress granule (SG) formation. The ectopic expression of G3BP1 in N-expressing cells was sufficient to reverse this phenotype. Since N is an RNA-binding protein, we performed iCLIP-sequencing experiments in cells, with or without exposure to oxidative stress, to identify the host RNAs targeted by N. Our results indicate that SARS-CoV-2 N protein binds directly to thousands of mRNAs under both conditions. Like the G3BPs stress granule proteins, N was found to predominantly bind its target mRNAs in their 3’UTRs. RNA sequencing experiments indicated that expression of N results in wide-spread gene expression changes in both unstressed and oxidatively stressed cells. We suggest that N regulates host gene expression by both attenuating stress granules and binding directly to target mRNAs.

## Introduction

Severe acute respiratory syndrome coronavirus 2 (SARS-CoV-2), the causative agent of the ongoing global pandemic of ‘Coronavirus Disease 2019 (COVID-19), is an enveloped, positive-sense, single stranded RNA virus^1^. The SARS-CoV-2 virion includes genomic RNA (~30 kb) and four structural proteins, the crownlike spike (S) glycoprotein that binds to human ACE2 as a receptor^2^, the membrane (M) protein responsible for viral assembly in the endoplasmic reticulum, the ion channel envelope (E) protein, and the nucleocapsid protein (N) that assembles with the viral RNA to form the nucleocapsid^13^.

The N protein is a multifunctional RNA-binding protein that is involved in several aspects of the viral life cycle, including viral genomic RNA replication and virion assembly^4–6^. Furthermore, the N protein is produced at high levels in infected cells^4^, where it appears to enhance the efficiency of sub-genomic viral RNA transcription and modulate host cell metabolism^reviewed by 5^. The SARS-CoV-2 N protein has two distinct RNA-binding domains, an N-terminal domain [NTD] and a C-terminal domain [CTD], connected by a linker region (LKR) containing a serine/arginine-rich (SR-rich) domain (SRD)^7–9^. Several critical residues of the N protein have been shown to bind viral genomic RNA and modulate infectivity^7^. Recent studies also show that SARS-Cov-2 N protein undergoes RNA-induced liquid-liquid phase separation (LLPS)^10,11^. The LLPS of N promotes the cooperative association of the RNA-dependent RNA polymerase complex with viral RNA, thus maximizing viral replication^10^.

Recent proteomic studies have indicated that SARS-CoV-2 N associates with the host stress granule (SG)-nucleating proteins Ras-GTPase-activating protein SH3-domain-binding protein 1 and 2 (G3BP1 and G3BP2)^12–14^. SGs are membrane-less protein-mRNA aggregates that form in the cytoplasm in response to a variety of environmental stressors, such as oxidative stress, osmotic stress, UV irradiation, and viral infection^15,16^. Assembly of SGs correlates with the arrest of translation initiation followed by polysome disassembly, resulting in an increase of uncoated mRNAs in the cytoplasm. Therefore, SGs are thought to protect endogenous mRNA from stress-mediated degradation. G3BP1 and G3BP2 are considered ‘essential’ for SG formation^17,18^, and overexpression of these proteins results in SG formation even in the absence of stress^19^. Recent studies have identified many RNA-binding proteins (RBPs), including G3BP1/2, TIA1, PRRC2C and UBAP2L, that appear to be essential for SG formation via LLPS^20^. SGs are generally believed to have an antiviral role upon viral infection^16^, and many viruses manipulate SGs to evade host responses^21^. For example, it has been found that the Zika virus hijacks key SG proteins, including G3BP1 and CAPRIN-1, via an interaction with the capsid^22^. This inhibits the formation of SGs and benefits viral replication^22^. Currently, however, the functional significance of SARS-CoV-2 N protein’s interaction with G3BPs remains unknown^23^.

Considering its critical roles throughout viral infection, the SARS-CoV-2 N protein has been described as a promising target for vaccine and antiviral drug development^4,24^. Recently, it has been observed that, upon expression of the SARS-CoV-2 N protein in human induced pluripotent stem cells (iPSC), pluripotency is abolished and the proliferation rate reduced^25^. Furthermore, it was found that long term N expression drives iPSC to fibroblasts^25^. Despite the urgent need to investigate molecular mechanisms through which N protein affects infected cells, there has been a lack of information regarding its impact on the host metabolism and transcriptome.

Here we show that the SARS-CoV-2 N protein physically interacts with G3BP1/2 and attenuates SG formation. We found that, like stress granule proteins G3BP1/2, N protein binds directly to cellular mRNAs, with a preference for 3’UTRs. Furthermore, we report that expression of N alters the expression profile of genes implicated in major biological processes and cellular pathways. We suggest that N protein alters host gene expression levels via both SG attenuation and direct interaction with mRNAs of many genes.

## Results

### Stress granule proteins G3BP1 and G3BP2 specifically co-purify with the SARS-CoV-2 N protein

To examine the potential impacts of viral proteins on host cell metabolism, we generated HEK293 cell lines expressing EGFP-tagged variants of 21 full-length SARS-Cov-2 genes (Supplemental Table S1). Each of the EGFP-tagged proteins was subjected to affinity purification with anti-GFP antibodies followed by Orbitrap-based precision mass spectrometry analysis (AP-MS). To remove any possible indirect associations mediated by DNA or RNA, cell extracts were treated with a promiscuous nuclease (Benzonase) prior to the affinity pull-downs. We scored specific protein interactions against GFP control purifications using SAINTexpress analysis^26^ with a statistical cutoff of ≤ 1% false discovery rate [FDR]. In total, we identified 647 protein-protein interactions (PPIs) involving 277 unique cellular proteins (Extended Figure 1; Supplemental Table S1). We performed KEGG enrichment analysis using the purified interaction partners and identified major cellular pathways, including RNA transport, protein processing in the endoplasmic reticulum, the proteosome and oxidative phosphorylation (P-value cut-off 0.05 (FDR)) (Extended Figure 2A, B). In addition, Huntington, Parkinson’s and Alzheimer’s disease related pathways were significantly enriched among the identified interaction partners (P-value cut-off 0.05 (FDR)) (Extended Figure 2B). We next compared our protein interaction data with two previously published AP-MS studies that were also performed in HEK293 cells^13,14^. We observed that there is little interaction partner overlap among the different reports, highlighting the variability of the experimental and/or statistical analyses employed in the various studies to catalogue the virus-host PPIs (see discussion) (Figure 1A). Remarkably, the only shared interaction partners among all three studies were the stress granule nucleating proteins G3BP1 and G3BP2. Considering their essential role in stress granule formation^17,18^, we next focused our attention on examining the functional significance of G3BP1 and G3BP2 interaction with the SARS-CoV-2 proteins.

**Figure 1:**
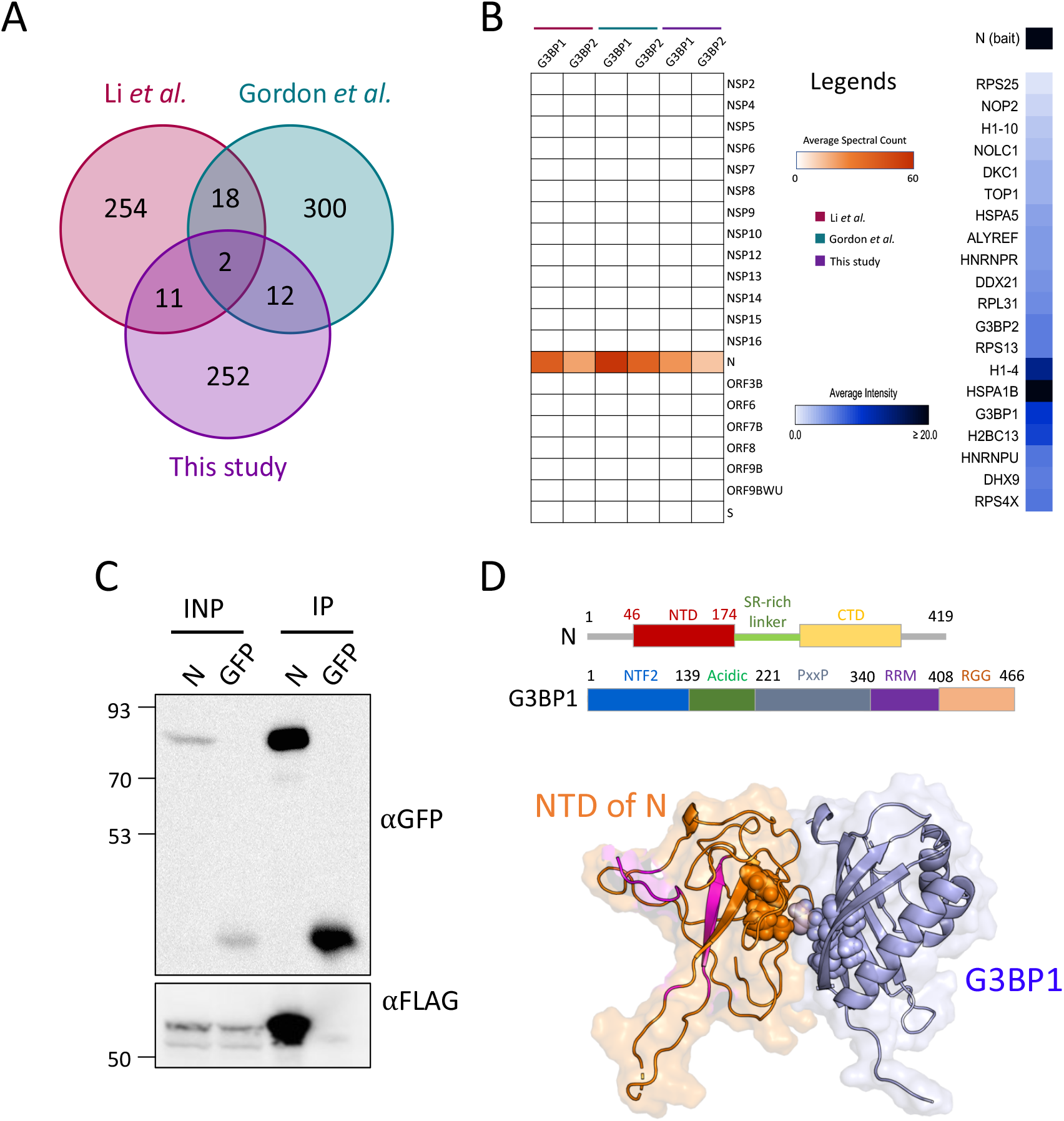
Stress granule proteins G3BP1 and G3BP2 interact with SARS-CoV-2 N protein. **A:** Venn diagram showing the overlap among three different AP-MS studies. Processed AP-MS data for Li et al. (PMID: 3283836) and Gordon et al. (PMID: 32353859) was downloaded from the supplemental files as published with each study. **B:** Left-Heatmap illustration of SARS-CoV-2 proteins’ interaction with G3BP1 and G3BP2 across three different studies. G3BP1 and G3BP2 co-purify with N protein. Right: Heatmap depiction of SARS-CoV-2 N protein-protein interactions as detected in this study (FDR≤1%). Note: Legends for each heatmap are provided in the middle panel. **C:** Interaction of SARS-CoV-2 N protein with G3BP1 detected by co-immunoprecipitation experiments. IP were performed with GFP antibodies using whole cell lysates prepared from HEK293 cells expressing both GFP-N (GEFP: 30kDa+N:45kDa) and FLAG-G3BP1 proteins. Cell lysates were treated with nuclease prior to IPs. Blots were probed with the indicated antibodies. **D: Top-** Domain organization of the N and G3BP1 proteins. **Bottom**-Cartoon illustration of protein docking studies. The complex structure model between G3BP1 (light blue cartoon) and the N protein (orange cartoon) is shown. The orange spheres and the light blue spheres are the predicted interface residues for N and G3BP1, respectively. The reported theoretical RNA-binding residues on N are highlighted in magenta.

Our AP-MS analysis indicated that, among the 21 examined SARS-CoV-2 proteins, G3BP1/2 co-purified exclusively with the Nucleocapsid N protein (Figure 1B; Extended Figure 1). We confirmed the interaction between N and G3BP1 by co-immunoprecipitation (co-IP) experiments using whole cell extracts (WCEs) prepared from cells expressing GFP-N and FLAG-G3BP1 (Figure 1C). Furthermore, the interaction was resistant to nuclease treatment indicating that it is not mediated by RNA (Figure 1C). The N protein NTD, which contains a pocket for binding viral RNA, is highly conserved across the coronaviruses and has been described as a potentially druggable target^8^. Recently, NMR structural studies identified several arginine (R) residues, including R92, R107, and R149, that directly contact the RNA^27^. Since crystal structures for both the SARS-CoV-2 N protein NTD and G3BP1 are available^27,28^, we performed protein docking studies to identify the residues on the two proteins that might come directly into contact. Our analyses predict a highly stable N-G3BP1 complex structure (Figure 1D), further supporting our AP-MS and co-IP analyses. We found that residues 90-96 (magenta spheres in Figure 1D) of G3BP1 (blue cartoon) potentially interact with residues 130-134 (green spheres) of the N protein (red cartoon) (Figure 1D). We conclude, therefore, that the SG resident proteins G3BP1/2 interact physically with the viral N protein and that this interaction is highly specific, as no other SARS-CoV-2 protein was observed to pull down G3BPs.

### Expression of N reduces stress granule formation

To begin elucidating the role of N in the context of SG formation, we performed AP-MS analysis using G3BP1 as the bait. SAINTexpress analysis indicated that, in addition to G3BP2, several SG nucleator proteins, including CAPRIN1, USP10, ATXN2l and NUFIP2, co-purify with G3BP1 as high-confidence (FDR≤0.01) interaction partners (Figure 2A). These high-confidence copurifying proteins were previously shown to function in SG formation upon exposure to stress^18,20,29^. Notably, the so-called ‘essential’ SG resident proteins that we identified as G3BP1 interaction partners were specifically depleted from the SARS-CoV-2 N interactome (compare Figures 1B and 2A). While further studies are underway to directly examine any remodelling of the G3BP1/2 PPIs upon N expression, our current analysis suggests that N might function to sequester G3BP1/2 away from interacting SG nucleating proteins, thus attenuating the SG formation.

**Figure 2:**
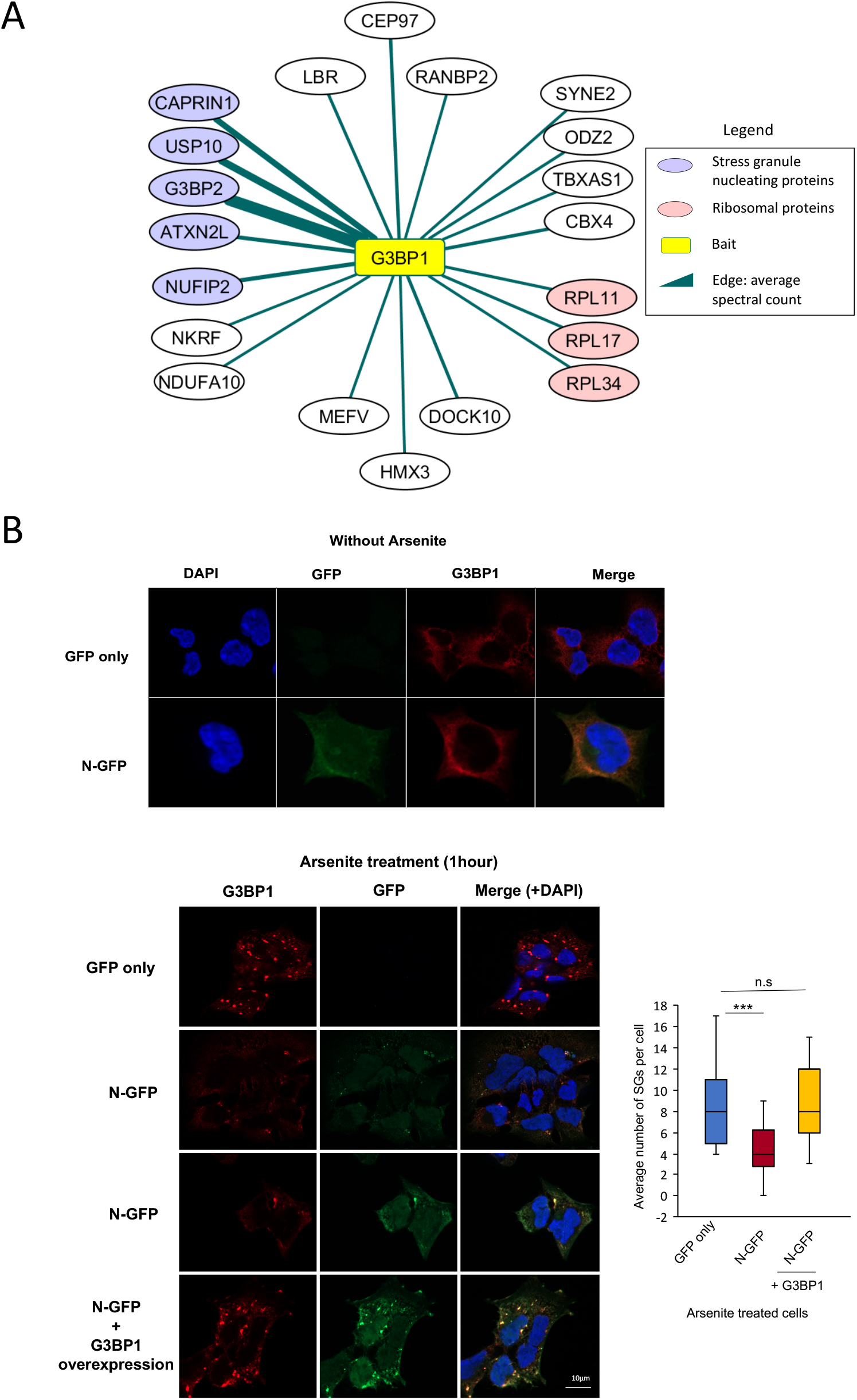
SARS-CoV-2 N protein attenuates stress granule formation. **A:** Network representation of high confidence (FDR≤0.01) G3BP1 co-purifying proteins. A figure legend is provided. **B:** Top panels-Immunofluorescence (IF) analysis to examine the localization of N and G3BP1 in HEK293 cells without NaAsO_2_ treatment. Bottom panels: IF was performed in HEK293 cells treated with NaAsO_2_ to quantify stress granule formation in GFP-N-expressing vs GFP only cells. Stress granules were identified by staining with anti-G3BP1 antibody. The box plot shows average number of stress granules per cell. Experiment was performed in biological replicate (n=2, total cells 50) and a student’s t test was used to examine the statistical significance (***p≤0.001). For nuclear counterstaining DAPI was used.

To test the hypothesis that N attenuates SG formation, we performed immunofluorescence studies. Cells expressing either GFP-N or GFP alone were subjected to oxidative stress by sodium arsenite (NaAsO_2_) treatment for one hour. As identified by G3BP1 staining (Figure 2B), SG punctae were readily observable in cells expressing both GFP and GFP-N Remarkably, we observed that the expression of N significantly reduced the number of observable SGs in cells subjected to NaAsO_2_ treatment (Figure 2B). Furthermore, we also observed that the N protein itself localized to the remaining SGs (Figure 2B). These data suggest that the expression of N negatively impacts SG formation upon exposure to stress.

Next, we examined whether this phenotype could be rescued through the overexpression of G3BP1 in N-expressing cells. N-expressing cells were transiently transfected with G3BP1 constructs followed by the induction of oxidative stress using NaAsO_2_. Consistent with the idea that N sequesters away G3BP1 from its interacting SG nucleators, we observed that the overexpression of G3BP1 rescued the SG attenuation phenotype (Figure 2B). As expected, N localized to the SGs in G3BP1 overexpressing cells. Overall, we conclude that N expression inhibits the formation of SGs in the host cells, possibly by sequestering away G3BPs from their interacting SG nucleating proteins.

### SARS-CoV-2 N protein directly binds host mRNAs with a preference for 3’UTRs

Since N protein contains RNA-binding domains^8^, we also examined whether N interacts directly with host mRNAs. We initially performed crosslinking and immunoprecipitation (CLIP) followed by gel electrophoresis and autoradiography to detect crosslinked RNA. We observed that N cross-links robustly to RNA *in vivo* upon exposure to UV (Figure 3A). In contrast, GFP alone did not yield any observable radioactive signal, indicating that N directly binds host RNAs (Figure 3A). Next, to identify host mRNAs bound by N, we carried out individual-nucleotide resolution UV crosslinking and immunoprecipitation (iCLIP) followed by high throughput sequencing (iCLIP-seq) experiments in biological replicates along with size-matched inputs (SMI) as controls.

**Figure 3:**
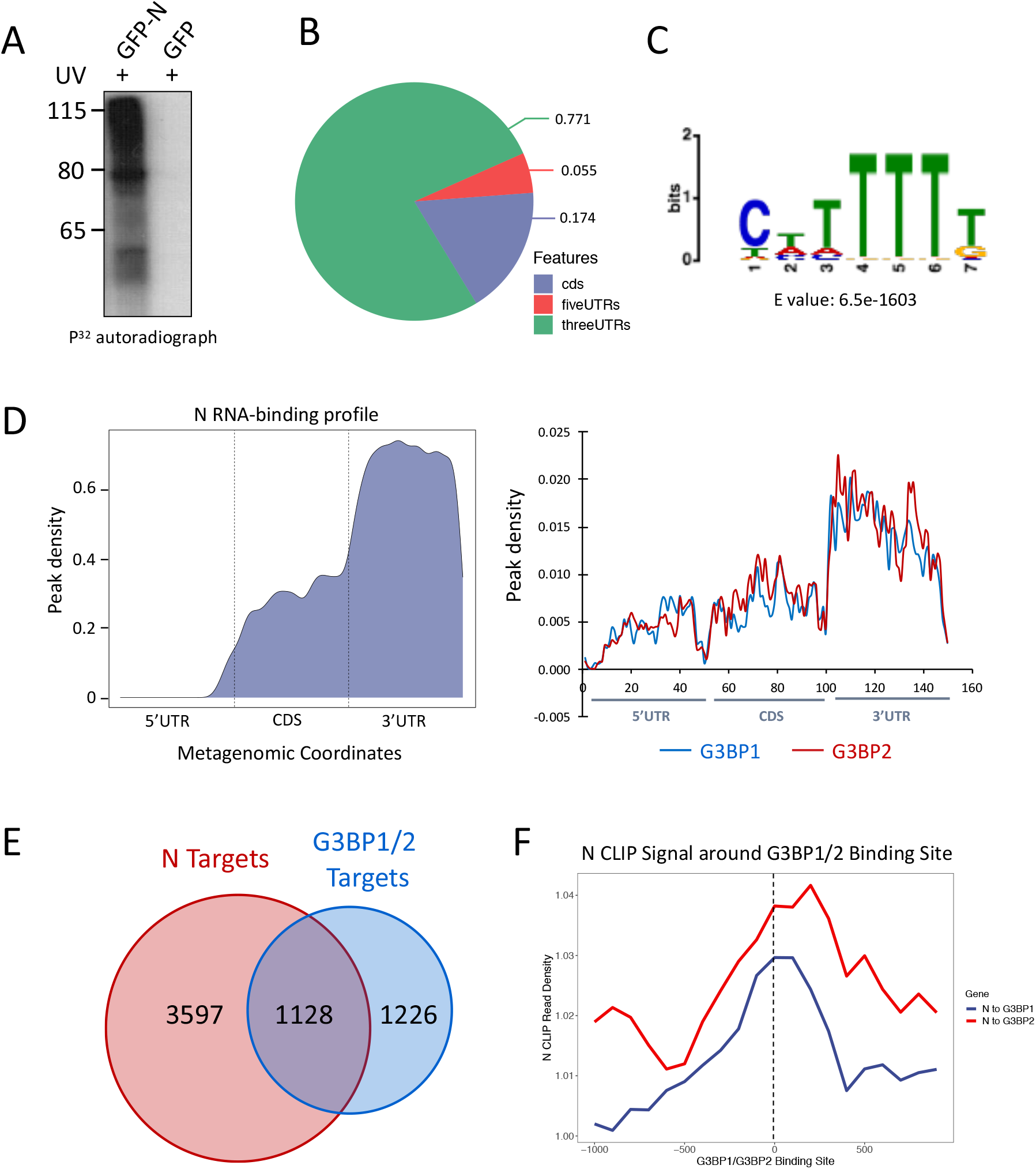
SARS-CoV-2 N directly binds to host mRNAs. **A:** Autoradiographs of immunopurified ^32^P-labeled N-RNA complexes after partial RNase I digestion. HEK293 cells were UV-crosslinked and GFP-N was immunoprecipitated using anti-GFP antibody. Purified RNA-protein complexes were resolved on 4-12% Bis-Tris gels after radiolabeling the RNA and transferred to nitrocellulose membranes. Cells expressing only GFP were used as negative control. **B:** Pie chart representing the distribution of N-bound RNAs. The majority of the N targets were found to be mRNAs. **C:** Enriched sequence motif found in N iCLIP-seq peaks. E-value represents the significance of the motif against control sequences. **D:** left-Standardized metaplot profile showing the normalized peak density of N-iCLIP. CDS represents the coding sequence. Right, Standardized metaplot profile showing the normalized peak density of G3BP1 and G3BP2. **E:** Venn diagram representing the overlap of targets between N and shared targets of G3BP1/2. **F:** Average iCLIP peak density of N around G3BP1 and G3BP2 binding sites. Note: To generate G3BP1 and G3BP2 metaplots and the Venn diagram, processed PAR-CLIP-seq data was used essentially as reported in the supplemental data set 3 of^30^.

Through peak calling in comparison with the SMI controls, we identified >30,000 high confidence peaks encompassing ~4500 unique human protein-coding genes (Extended Figure 3A; Supplemental Table S2). Gene ontology (GO) enrichment analysis was carried out using these N-bound human genes. Major biological processes related to post-transcriptional regulation, including mRNA processing, RNA catabolic processes, RNA transport, post-transcription regulation of gene expression, and translation regulation were significantly enriched (FDR 0.05) (Extended Figure 3B). KEGG enrichment analysis indicated that N-bound genes were enriched for pathways that included protein processing in endoplasmic reticulum, ribosome and cell cycle (FDR 0.05) (Extended Figure 3B).

We next examined the iCLIP peak distribution across the N-bound transcripts. Remarkably, most peaks (77% of the total peaks) were found within the annotated 3’ untranslated regions (UTRs) (Figure 3B). We also searched for enriched sequence motifs and found that U-rich sequence motifs were significantly enriched among the N-binding sites (Figure 3C; Extended Figure 3C). Since N interacted with G3BP1 and G3BP2, we included their PAR-CLIP-seq data^30^ for comparison. Our metagene analyses indicated that, similar to the RNA-binding profiles of G3BP1 and G3BP2, the SARS-CoV-2 N protein predominantly binds within the 3’UTRs of its target genes, consistent with its peak distribution (Figure 3D). Our analysis revealed that ~47% of the G3BP1 and G3BP2 targets were also bound by the N protein (Figure 3E). Furthermore, N’s iCLIP-seq signal was enriched around the G3BP1 and G3BP2 binding sites on the RNAs (Figure 3F). These results reinforce the idea that SARS-CoV-2 N and G3BPs might function together in the infected cells. It is conceivable that N might reshape the G3BP1/2-bound transcriptome to the advantage of SARS-CoV-2. Further experiments are underway to test this possibility (experimental data being analyzed). Additionally, studies are ongoing to investigate whether N binds cooperatively in conjunction with G3BPs on shared target mRNAs.

### SARS-CoV-2 N alters the host gene expression profile

To examine the effect of SARS-CoV-2 N on host gene expression, we performed RNA-seq analysis in HEK293 cells expressing GFP-N. Since N appears to have a role in SG formation, we also included RNA samples prepared from cells treated with NaAsO_2_. Cells expressing GFP alone were used as controls in these experiments. The RNA-seq replicates highly correlated with each other, indicating the reproducibility of our data (Extended Figure 4). Differential expression analysis identified 4363 and 2942 genes that were significantly differentially expressed in N-expressing cells that were untreated or NaAsO_2_-treated (Q<0.05), respectively (Figure 4A). Of the 4363 differential genes in the untreated samples 2207 were upregulated, whereas 2156 were downregulated (Supplemental data S3,4). In the NaAsO_2_-treated cells, 1765 and 1178 genes were up- and down-regulated, respectively. We found that ~82% of the differential genes in the NaAsO_2_ samples (2403/2942 genes) were the same as those that were also differentially expressed in the untreated cells. These results indicated that N affects a core set of genes under both conditions. Furthermore, after normalizing the NaAsO_2_ RNA-seq data against the untreated samples, we identified a small subset of genes significantly upregulated in the N-expressing cells under stress conditions (Q<0.05) (Supplemental data S5).

**Figure 4:**
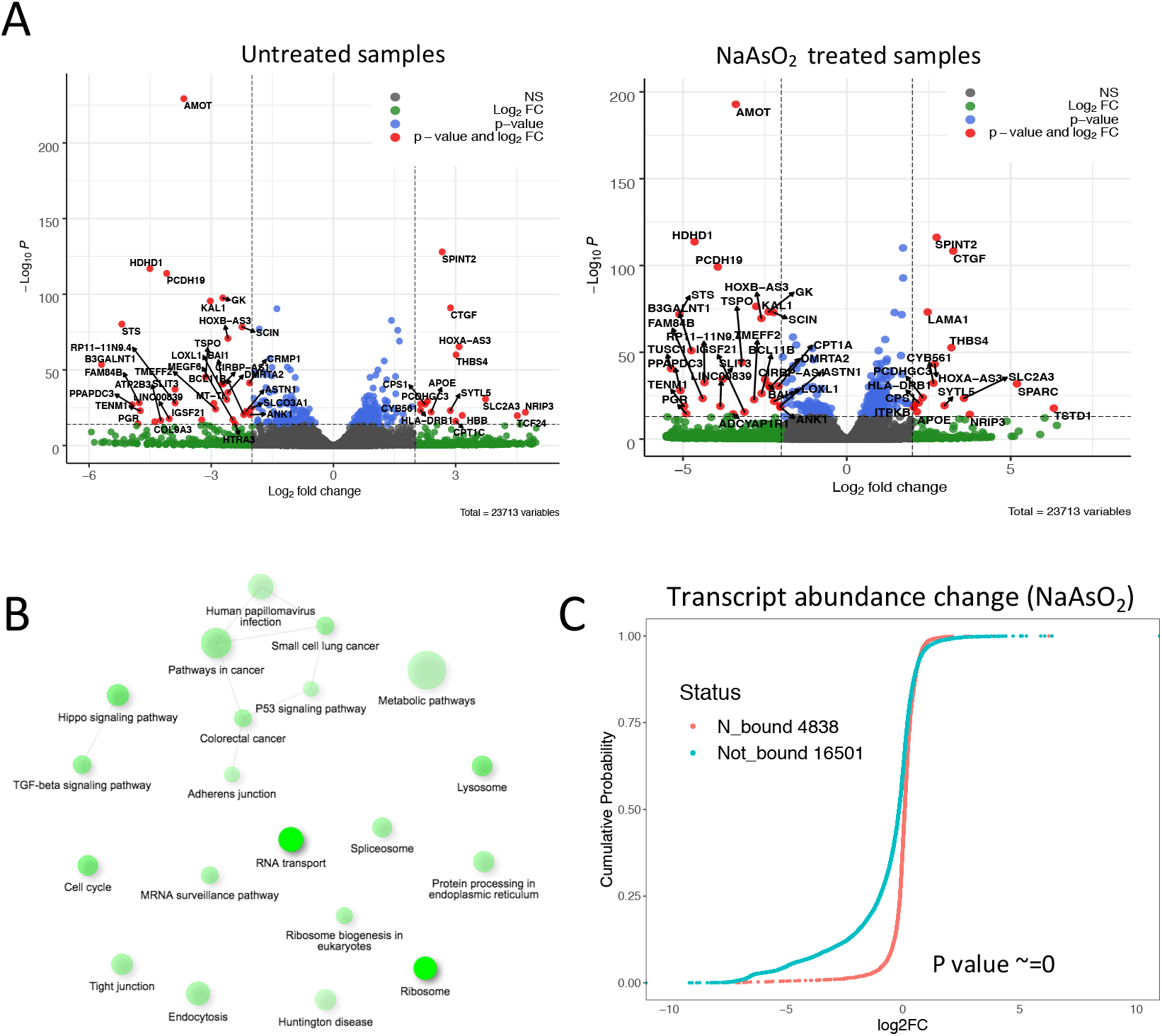
Expression of N results in alterations of host gene expression. **A:** Volcano plot representation of differentially expressed genes in untreated (left) and NaAsO_2_ treated cells (right). Differential expression was calculated against GFP expressing cells in both conditions. Each dot represents a single gene. Genes with Q≤0.05 were considered significant. Horizontal dotted lines represent log2fold change of 1 and highly significant genes are shown as red dots with label indicating the gene name. Grey dots represent genes that are considered as insignificant in this representation. Legends are provided at the top right corner of each plot. **B:** KEGG pathway enrichment analysis using genes that were significantly upregulated in N cells in comparison to the GFP cells. Only the top 20 most significant terms are shown (Q≤0.05). Darker nodes are more significantly enriched gene sets. Bigger nodes represent larger gene sets. Thicker edges represent more overlapped genes. **C:** Cumulative distribution analysis in abundance of N iCLIP target mRNAs after N overexpression in NaAsO_2_ treated HEK293 cells. Legend is provided on the plot.

We next examined whether the N-affected host genes are enriched for any specific biological processes. GO enrichment analysis indicated that the upregulated genes were significantly enriched in biological processes related to cellular localization, intracellular protein transport, and cell cycle regulation (Q-value cut-off 0.05; Extended Figure 5A). Furthermore, KEGG pathway analysis showed that cancer-related pathways, TGF-beta signalling, Hippo signalling, protein processing in ER and RNA transport were significantly enriched in the upregulated genes (Q-value cut-off 0.05) (Figure 4B). The downregulated genes, on the other hand, were enriched in pathways related to neurogenesis and nervous system development (Q-value cut-off 0.05; Extended Figure 5B). These data suggest that the expression of N results in deregulation of many functionally important genes.

Considering that N interacts with G3BP1, which has been shown to enhance the stability and translation of its target mRNAs^15,30,31^, we correlated the effect of N binding to its target mRNA abundance. Our results indicate that mRNAs associated with N were generally stabilized in comparison to the non-targets, which were significantly decreased in their abundance upon NaAsO_2_ treatment (Figure 4B). G3BP1 and G3BP2 also showed similar trends in our analyses (Extended Figure 6), consistent with their shared targets with N. These data suggest that the RNA-binding of N might stabilize certain target mRNAs against the stress-induced degradation (see discussion). Further studies are underway to directly identify the N targets in NaAsO_2_ treated cells and examine how N might reshape the G3BP1-bound transcriptome.

## Discussion

Given that the ongoing COVID-19 pandemic has globally caused more than one million deaths (World health organization data; October 5, 2020), there is an urgent need to better understand the SARS-CoV-2 life cycle for effective antiviral drug development. In this study, we report the biological impact of the SARS-CoV-2 N protein on host cells. Starting from a proteomic workflow, we found that there is little overlap across studies that recently reported virus-host PPIs for SARS-CoV-2 proteins^13,14^. This observation is perhaps not surprising due to varied experimental and statistical analyses employed across different studies. We suggest that, in the absence of additional evidence to support these conclusions, caution should be exercised in interpreting proteomics data for drug development

Among the high-confidence interactors detected across three AP-MS studies were the stress granule resident proteins G3BP1 and G3BP2. SGs are typically formed upon viral infection, possibly as a cellular response to block viral replication^16^. However, viruses have evolved many counter-measures to combat host responses to viral infection and, indeed, many viruses have been reported to destabilize SGs upon infection^15,16^. Certain viruses have been reported to hijack G3BP1 and G3BP2, inhibiting SG formation to the benefit of the virus. In such cases, depletion of G3BPs (or other SG components) reduced viral replication^15,16^. For example, Zika virus hijacks G3BPs, to reduce SG formation and benefit viral replication^22^, and it was shown that depletion of G3BP1 indeed reduced Zika virus replication^22^. In contrast to Zika, however, G3BPs have been found to inhibit the replication of Sendai virus and vesicular stomatitis virus^32^. Here we showed that SARS-CoV-2 N protein interacts physically with G3BPs and that expression of N attenuates SG formation. It remains to be seen, however, whether or not G3BPs also play a role in SARS-CoV-2 replication. In this context, it is interesting to note that a recent study showed that G3BPs interact with SARS-CoV-2 RNA^33^.

Although N expression resulted in both up- and down-regulation of various host genes, N-bound mRNAs were more stable in comparison to non-targets. Previous reports have shown that certain adenovirus and hepatitis C virus (HCV) proteins stabilize host mRNAs via binding to AU-rich elements present within 3’UTRs of target transcripts^34,35^. AU-rich elements (ARE), which are present in many proto-oncogenes, growth factor and cytokine mRNAs, target mRNAs for degradation^36–38^. The adenovirus protein, E4orf6, has been reported to stabilize the host ARE-containing mRNAs, and this stabilization was found to be necessary for its oncogenic activity^34^.

Similarly, NS5A of HCV was shown to directly bind to U- (or G-) rich elements within 3’UTRs of host mRNAs, resulting in their stabilization^35^. The NS5A target mRNAs were found to be highly enriched in regulators of cell growth, cell death and cancer^35^. Based on these findings, it has been suggested that virus-induced stabilization of host transcripts prevents cell death and promotes growth of virus-infected cells^35^. While SARS-CoV-2 is an acute respiratory virus, our finding that N protein recognises U-rich elements and may stabilize its targets via directly binding to 3’UTRs of host mRNAs is of particular interest given that the up-regulated genes were significantly enriched in cancer-related pathways. Although mechanistic details of N-mediated stabilization remain unknown, we found that N and G3BPs not only physically interact with each other but also have similar RNA-binding profiles with many shared targets. G3BPs have been shown to stabilize their target mRNAs in unstressed as well as stressed cells^reviewed by 15^. We suggest that SARS-CoV-2 N and G3BPs bind to various target mRNAs, resulting in their stabilization.

It is worth noting that N-bound stabilized targets were also enriched in major cellular pathways, including “protein processing in ER” and “TGF-beta signalling”. This finding is of interest, as viruses, including SARS-CoV, have been shown to cause ER stress and the induction of signalling pathways collectively known as the unfolded protein response (UPR)^39,40^. However, coronaviruses are thought to have evolved the ability to subvert, or even exploit, certain aspects of the UPR and overcome protein translation shutdown^39^. Further studies will be required to investigate whether or not the stabilization and/or direct binding of N to ER-related mRNAs plays a role in the ER restructuring that is observed upon infection.

Further studies are underway to investigate the role of N upon oxidative stress, including how N protein might rewire the G3BP RNA-binding profile and what mechanisms N protein might use to reshape the host transcriptome. Based on the evidence presented here, we suggest that SARS-CoV-2 N protein affects host cell metabolism and gene expression via multiple pathways, including SG attenuation, sequestration of G3BPs, and direct binding to mRNAs of some functionally important genes. Considering that N is a promising target for drug and vaccine development, our findings might be of therapeutic interest.

## Materials and Methods

### Cell cultures

HEK293 cells (Flp-In 293 T-REx cell lines) were obtained from Life Technologies (Invitrogen catalogue number R780-07). Cell cultures were maintained in Dulbecco’s modified Eagle’s medium (DMEM) (Wisent Bioproducts catalogue number 319-005-CL), which was supplemented with 10% FBS (Wisent Bioproducts catalogue number 080-705), sodium pyruvate, non-essential amino acids, and penicillin/streptomycin as described^41^.

### Epitope tagging in HEK293 cells

The gateway-compatible entry clones for 21 viral ORFs^42^ (kindly provided by Dr. Frederick P. Roth) were cloned into the pDEST pcDNA5/FRT/TO-eGFP vector according to the manufacturer’s instructions to create fusions between eGFP and viral proteins. The vectors were co-transfected into Flp-In T-REx 293 cells together with the pOG44 Flp recombinase expression plasmid. Cells were selected for FRT site-specific recombination into the genome, and stable FlpIn 293 T-REx cell lines were maintained with hygromycin (Life Technologies, 10687010) at 2ug/ml. Doxycycline was added to the culture medium 24 h before harvesting to induce the expression of the various viral genes of interest.

### Antibodies

The following antibodies were used in this work: Flag (Sigma monoclonal antibody catalogue number F1804); GFP (Abcam polyclonal antibody catalogue number 290); GFP monoclonal antibody (Life Technologies G10362); G3BP1 (Santa Cruz monoclonal antibody catalogue number sc-365338).

### Immunoprecipitation (IP) and western blots

To perform Co-IP experiments, cell pellets were lysed in 1mL of lysis buffer (140 mM NaCl, 10 mM Tris pH 7.6–8.0, 1% Triton X-100, 0.1% sodium deoxy-cholate, 1 mM EDTA) containing protease inhibitors (Roche catalogue number 05892791001). Cell extracts were incubated with 75 units of Benzonase (Sigma E1014) for 30min in a cold room with end-to-end rotation. The cell lysates were cleared in a microcentrifuge at 15,000 g for 30 minutes at 4°C. The supernatant was transferred to a new tube and incubated with 1μg of GFP antibody for 4 hours to overnight, and subsequently 10 μl protein G Dyna beads were added and incubated for an additional 2 hours (Invitrogen catalogue number 10003D). The samples were washed three times with lysis buffer containing an additional 2% NP40 for 5 min each in a cold room with end-to-end rotation. The samples were then boiled in SDS gel sample buffer. Samples were resolved using 4-12% BisTris-PAGE and transferred to a PVDF membranes (Bio-Rad catalogue number 162-0177) using a Gel Transfer Cell (BioRad catalogue number 1703930). Primary antibodies were used at 1: 5000 dilution, and horseradish peroxidase-conjugated goat anti-mouse (Thermo Fisher 31430) or antirabbit secondary (Thermo Fisher 31460) antibodies were used at 1:10,000. Blots were developed using Pierce ECL Western Blotting Substrate (Thermo Scientific catalogue numbers 32106).

### iCLIP-seq experiments

Individual nucleotide resolution UV crosslinking and immunoprecipitation (iCLIP) was performed as previously described^43^ with the modifications as detailed in our previous report^44^. Briefly, cells were grown in 25cm culture plates and were UV cross-linked with 0.4 J/cm^2^ at 254 nm in a Stratalinker 1800 after induction with Doxycycline for 24 hours. Cells were lysed in 2mL of iCLIP lysis buffer. 1mL of lysate was incubated with 2 μl Turbo DNase (Life Technologies catalogue number AM2238) and RNase I (1:250; Ambion catalogue number AM2294) for 5 min at 37°C to digest the genomic DNA and obtain RNA fragments of an optimal size range. GFP-N was immunoprecipitated from two independent HEK293 cell pellets using 5μg of anti-GFP antibody (Life Technologies G10362). A total of 2% input material was obtained for size-matched control libraries (SMI) prior to the IPs. Following stringent washes with iCLIP high salt buffer and dephosphorylation with T4 PNK, on-bead-ligation of pre-adenylated adaptors to the 3’-ends of RNAs was performed using the enhanced CLIP ligation method. The immunoprecipitated RNA was 5’-end-labeled with ^32^P using T4 polynucleotide kinase (New England Biolabs catalogue number M0201L), separated using 4–12% BisTris-PAGE and transferred to a nitrocellulose membrane (Protran). For the input sample, the membrane was cut matching the size of the IP material. RNA was recovered by digesting proteins using proteinase K (Thermo Fisher catalogue number 25530049) and subsequently reverse transcribed into cDNA. The cDNA was size-selected (low: 70 to 85 nt, middle: 85 to 110 nt, and high: 110 to 180 nt), circularized to add the adaptor to the 5’-end, linearized, and then PCR amplified using AccuPrime SuperMix I (Thermo Fisher catalog number 12344040). The final PCR libraries were purified on PCR purification columns (QIAGEN), and the eluted DNA was mixed at a ratio of 1:5:5 from the low, middle, and high fractions and submitted for sequencing on an Illumina HiSeq2500 to generate single-end 51 nucleotide reads with 40M read depth.

The barcoded primers used were:

N iCLIP replicate 1: Rt9clip
/5Phos/NNGCCANNNAGATCGGAAGAGCGTCGTGgatcCTGAACCGC
N iCLIP replicate 2: Rt16clip
/5Phos/NNTTAANNNAGATCGGAAGAGCGTCGTGgatcCTGAACCGC
N size-matched Input: Rt10clip
/5Phos/NNGACCNNNAGATCGGAAGAGCGTCGTGgatcCTGAACCGC

### Affinity purification followed by mass spectrometry

The AP-MS procedure for HEK293 cells was performed essentially as previously described^41,45^. Briefly, ~20×10^6^ cells were grown in two independent batches representing biological replicates. After 24 h induction of protein expression using doxycycline, cells were harvested. Cell pellets were lysed in high-salt NP-40 lysis buffer (10 mM Tris-HCl pH 8.0, 420 mM NaCl, 0.1% NP-40, plus protease/phosphatase inhibitors) with three freeze-thaw cycles. The lysate was sonicated as described^41^. To remove genomic DNA and RNA, we treated cell lysates with Benzonase for 30 min at 4°C with end-to-end rotation. The WCE was centrifuged to pellet any cellular debris. GFP-tagged viral proteins were immunoprecipitated with anti-GFP antibody (G10362, Life Technologies) overnight followed by a 2-hour incubation with Protein G Dynabeads (Invitrogen). The beads were washed 3 times with buffer (10mM TRIS-HCl, pH7.9, 420mM NaCl, 0.1% NP-40) with end-to-end rotation in the cold room and twice with buffer without detergent (10mM TRIS-HCl, pH7.9, 420mM NaCl). The immunoprecipitated proteins were eluted with 0.5M NH_4_OH and lyophilized.

### Sample Preparation and Proteomic Analysis

Each purified elute was digested in-solution with trypsin for MS analysis. Briefly, each sample was resuspended in 44uL of 50mM NH_4_HCO_3_, reduced with 100mM TCEP-HCL, alkylated with 500mM iodoacetamide for 45 min in the dark room, and digested with 1ug of trypsin overnight at 37°C. Samples were desalted using ZipTip Pipette tips (EMD Millipore) using standard procedures. The desalted samples were analyzed with an LTQ-Orbitrap Velos mass spectrometer (ThermoFisher Scientific) utilizing a 90-minute HPLC gradient and top 15 data-dependent acquisition.

### Experimental design for mass spectrometry

We processed independently at least two biological replicates for each bait along with negative controls in each batch of sample. Material from HEK293 cells expressing GFP only was used as control. We performed extensive washes between samples to minimize carry-over. Furthermore, the order of sample acquisition on the mass spectrometer was reversed for the second replicate to avoid systematic bias.

### Mass-spectrometry Data Analysis

AP-MS datasets were searched with Maxquant (v.1.6.6.0)^46^. Human protein reference sequences from the UniProt Swiss-Prot database was downloaded on 18-06-2020. SARS-CoV-2 protein reference sequences (GenBank accession NC_045512.2, isolate=Wuhan-Hu-1) were downloaded from https://www.ncbi.nlm.nih.gov/datasets/coronavirus/proteins_on_14-09-2020, supplemented by isolate 2019-nCoV/USA-WA1/2020 (GenBank Accession MN985325) for ORF3b, ORF9b and ORF9c. Spectral counts as well as MS intensities for each identified protein were extracted from Maxquant protein Groups file. The resulting data were filtered using SAINTexpress^26^ to obtain confidence values utilizing our two biological replicates. SAINTexpress BFDR 0.01 was chosen as a cut-off. The AP-MS data generated from HEK293 cells expressing GFP alone were used as negative control for SAINTexpress analysis.

### MS data visualization, functional enrichment and archiving

We used Cytoscape (V3.4.0;^47^) to generate protein-protein interaction networks. For better illustration, individual nodes were manually arranged in physical complexes. Dot plots and heatmaps were generated using ProHits-viz^48^. Functional enrichment of PPI data was performed using ShinyGO (v0.61)^49^ which utilizes hypergeometric distribution followed by FDR correction (FDR cutoff was 0.05). All MS files used in this study were deposited at MassIVE (http://massive.ucsd.edu).

### Prediction of complex structure model between G3BP1 and N protein

The crystal structures for G3BP1 and the N protein were downloaded from the Protein Data Bank (PDB) to build their complex structure model (PDB ID: 4FCJ_A and 6M3M_B). The amino acid sequence for each protein was extracted from the corresponding structure files according to the ‘ATOM’ records. We generated MSA for each protein by running HHblits^50^ with 8 iterations and E-value=1E-20 to search through the Uniclust30 library. The G3BP1 had 9435 homologous sequences and the N protein had 2665. These sequences were paired based on genomic distance or phylogeny. However, only the origin sequence was left after this concatenation. Thus, we input the concatenated single sequence to the deep learning-based algorithm trRosettato^51^ to predict the interface residues. Based on the trRosetta prediction, we found the residues 90-96 (light blue spheres in Figure 1) of G3BP1 (blue cartoon) tend to interact with the residues 130-134 (orange spheres) of the N protein (orange cartoon). The above interface residues were used as distance constraints to build a complex structure model (Figure 1) with the HDOCK server^52^. Finally, we utilized the Rosetta docking protocol to optimize the interface with local refinement^53^. The interface energy of the final model was −5.46, which indicates a reliable model according to the Rosetta document^53^.

### Sodium Arsenite (NaAsO_2_) treatment

HEK293 cells were treated with 0.5mM sodium arsenite (NaAsO_2_) for one hour. Briefly, we used HEK293 cells expressing SARS-CoV-2 N protein tagged with GFP (GFP-N), GFP alone, and ‘GFP-N + untagged G3BP1’. The expression was induced with doxycycline (1μg/ml) for 24 hours prior to NaAsO_2_ treatment for one hour.

### Immunofluorescence

SARS-CoV-2 GFP-N and GFP expressing HEK293 cells were seeded on poly-L-lysine coated and acid-washed coverslips. The expression of the proteins was induced using 1μg/ml doxycycline for 24 hours. Sodium arsenite (NaAsO_2_) treatment was performed as described above for 60 minutes prior to cell fixation. NaAsO_2_ was removed and cells were washed three times with PBS. Cells were fixed in 4% Paraformaldeyde for 15 minutes. Cells were subsequently permeabilized with 0.2% Triton X-100 in PBS for 5 min and incubated with block solution (1% goat serum, 1% BSA, 0.5% Tween-20 in PBS) for 1 hour. Santa Cruz G3BP (H-10) antibody was used for staining at 1:100 concentration in block solution for 2 hours at room temperature (RT). Cells were incubated with Goat anti-mouse secondary antibody and Hoescht stain in block solution for 1 hour at room temperature. Cells were fixed in Dako Fluorescence Mounting Medium (S3023). Imaging was performed the next day using a Zeiss confocal spinning disc AxioObserverZ1 microscope equipped with an Axiocam 506 camera using Zen software. A single focal plane was imaged, and stress granule quantification was performed for each replicate (n=2; number of cells 50). The number of stress granules per cell was plotted as box plot, and statistical significance was calculated using the student’s t test.

### RNA extraction and sequencing

Total RNA was extracted using the RNeasy extraction kit (Qiagen) following the manufacturer’s instructions. Two independent biological samples for each condition were generated, resulting in a total of eight samples. DNase-treated total RNA was then quantified using Qubit RNA BR (cat # Q10211, Thermo Fisher Scientific Inc., Waltham, USA) fluorescent chemistry and 1 ng was used to obtain RNA Integrity Number (RIN) using the Bioanalyzer RNA 6000 Pico kit (cat # 5067-1513, Agilent Technologies Inc., Santa Clara, USA). Lowest RIN was 9.3; median RIN score was 9.75.

1000 ng per sample was then processed using the NEBNext Ultra II Directional RNA Library Prep Kit for Illumina (cat # E7760L; New England Biolabs, Ipswich, USA; protocol v. v3.1_5/20) including PolyA selection, with 15 minutes of fragmentation at 94 °C and 8 cycles of amplification. 1uL top stock of each purified final library was run on an Agilent Bioanalyzer dsDNA High Sensitivity chip (cat # 5067-4626, Agilent Technologies Inc., Santa Clara, USA). The libraries were quantified using the Quant-iT dsDNA high-sensitivity (cat # Q33120, Thermo Fisher Scientific Inc., Waltham, USA) and were pooled at equimolar ratios after size-adjustment. The final pool was run on an Agilent Bioanalyzer dsDNA High Sensitivity chip and quantified using NEBNext Library Quant Kit for Illumina (cat # E7630L, New England Biolabs, Ipswich, USA).”

The quantified pool was hybridized at a final concentration of 2.215 pM and sequenced single-end on the Illumina NextSeq 500 platform using a full High-Output v2.5 flowcell at 75 bp read lengths, for an average of 52 million pass-filter clusters per sample.

### Quantification and Statistical Analysis

#### Alignment and Read Processing

iCLIP libraries were demultiplexed using XXXNNNNXX barcodes, where X is a random nucleotide. The 5’barcode and the illumine adaptor was trimmed using Cutadapt^54^ (ver 2.10). The PCR duplicates were collapsed using UMI-tools^55^ (ver 1.0.1). RNA-seq and iCLIP library reads were mapped to Gencode assembly^56^ (GRCh37.p13) using STAR^57^ (ver 2.7.1). Only the uniquely mapping reads were used for the downstream analyses.

### RNA-seq Analysis

To identify the differentially expressed genes from RNA-seq data, we used DESeq2^58^ (ver 3.11) on gene counts generated using STAR and the human Gencode annotation V19. We filtered out genes with less than 10 counts across the sum of all RNA-seq samples. To plot differentially expressed genes as volcano plots, we used R-package EnhancedVolcanoplot (https://github.com/kevinblighe/EnhancedVolcano). The Gene Ontology enrichment analysis was performed and visualized using clusterProfiler^59^ (ver 3.16.1), with the universe set to all the genes detected by RNA-seq. Multiple testing correction was performed using the Benjamini-Hochberg method and q-value cut off of 0.05 was used. Additionally, GO/ KEGG enrichment was performed using ShinyGO (v0.61), which utilizes hypergeometric distribution followed by FDR correction where FDR cutoff was set to 0.05.

### iCLIP Metagene and Motif Analysis

Significant iCLIP peaks were called using Pureclip^60^, with input control and default settings, and the cross-linking sites within 50 bps from each other were merged. To visualize the distribution of N protein on genes, we used MetaPlotR^61^ to calculate and scale the distance of N protein peaks relative to transcriptomic features. The processed PAR-CLIP peaks for G3BP1 and G3BP2 were downloaded from GEO (accession code: GSE98 8 5 6)^32^. The read densities over ±1000 bp regions surrounding G3BP1 and G3BP2 peaks were calculated by diving the CLIP bpm over input bpm, where the bin size was set to 100. We used bedtools^62^ (ver 2.29.2) to retrieve DNA sequences from the iCLIP peaks and subjected the sequences to motif analysis using the MEME-ChIP suite^63^ (version 5.1.1).

## Acknowledgement

Authors would like to thank Dr. Frederick P. Roth for providing the SARS-CoV-2 ORFs. We also thank Ulrich Braunschweig for his help with CLIP data download. Tanja Durbic and Kyle Turner at the Donnelly Sequencing Centre are gratefully acknowledged for their assistance with next-generation sequencing.

## Authors’ contribution

SN-S and JFG conceived and designed the study. SN-S performed iCLIP-seq, participated in IFs, RNA-seq, Co-IP experiments, contributed to the computational analyses and co-ordinated the project. HL performed iCLIP and RNA-seq data analyses and participated in manuscript editing. NA performed Co-IPs, participated in all experimental work and manuscript preparation. EM performed AP-MS experiments. SF conducted IF and RNA extraction experiments. SP analyzed the AP-MS data. GB performed RT-qPCR experiments and edited manuscript. KA participated in iCLIP data analyses. HW conducted the docking studies. GZ and HT generated cell lines and performed AP-MS. JY supervised HW. BB supervised SF and edited the manuscript. ZZ coordinated computational analyses, supervised HL, and edited the manuscript. JFG supervised the project and data analyses. JFG and SN-S wrote the manuscript.

