## Supplemental Figures 1-6 for "SARS-CoV-2 Nucleocapsid protein attenuates stress granule formation and alters gene expression via direct interaction with host mRNAs"

**Extended Figure 1:** Dot plot overview of the interaction partners identified with 21 SARS-CoV-2 proteins expressed in HEK293 cells. Inner circle color represents the average spectral count, the circle size maps to the relative prey abundance across all samples shown, and the circle outer edge represents the SAINT FDR. Legend is provided.

**Extended Figure 2:** KEGG pathway (**A**) and Biological processes (**B**) enrichment analysis using cellular proteins identified as high confidence (FDR$\leq$1%) interaction partners for the SARS-CoV-2 proteins. Only top 10 most significant terms are shown (Q$\leq$0.05). Darker nodes are more significantly enriched gene sets. Bigger nodes represent larger gene sets. Thicker edges represent more overlapped genes.

**Extended Figure 3: A-** Pie chart representing the distribution of N iCLIP peaks across various RNA categories. Majority of the peaks were identified in the 3' untranslated region (3'UTR) of the mRNAs. **B-** GO enrichment analysis using N target genes as identified by iCLIP-seq. Dot plot on the left represents enrichment for biological processes and network on the right indicates KEGG pathways that were found to be enriched in the N targets. Legend is provided for the dot plot. **C:** Left- Five most significantly enriched sequence motifs as identified by the MEME software using N iCLIP peaks. E-values are provided below each motif to indicate the statistical significance over background sequences. Right- Centrino output plot indicating the distribution of the top two motifs around N's crosslinking sites on the RNAs as identified by iCLIP. Motifs were found to be centrally enriched.

**Extended Figure 4:** Principal component analysis showing the correlation between RNA-seq replicates for different conditions. Figure legend is provided.

**Extended Figure 5: A-** GO enrichment analysis using genes that were significantly upregulated in N cells in comparison to the GFP cells. Only top 10 most significant terms related to biological processes are shown (Q≤0.05). Darker nodes are more significantly enriched gene sets. Bigger nodes represent larger gene sets. Thicker edges represent more overlapped genes. **B-** GO enrichment analysis using genes that were significantly downregulated in N cells in comparison to the GFP cells. Figure legend same as described for A.

**Extended Figure 6:** Cumulative distribution analysis in abundance of G3BP1, G3BP2 and N shared iCLIP target mRNAs after N overexpression in HEK293 cells. Legend is provided.

Extended Figure 1

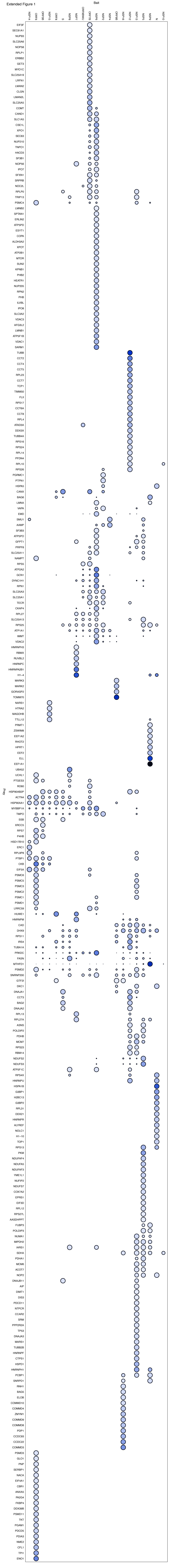

### Figure legend

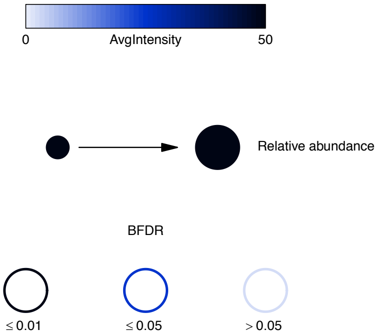

Extended Figure 2

A

**KEGG pathway enrichment**

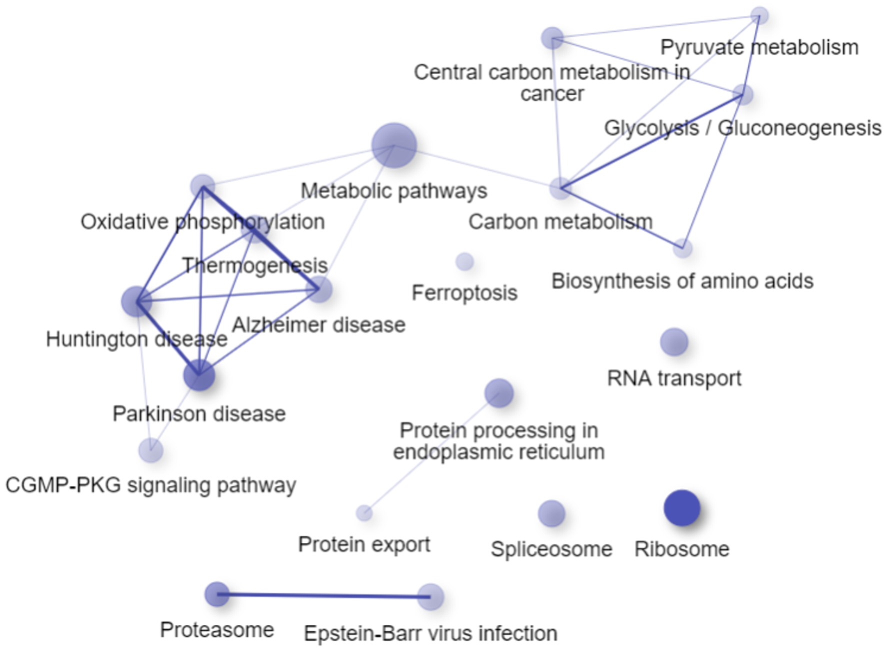


B

**Biological processes enrichment**

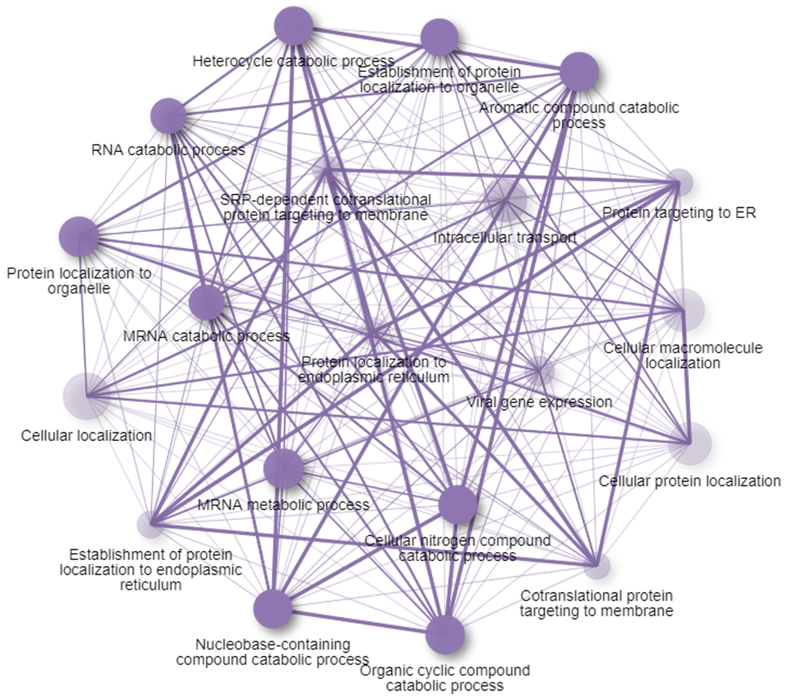

Extended Figure 3

A

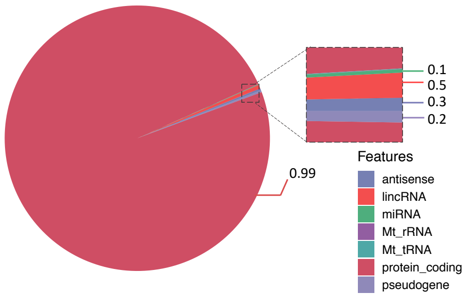


B

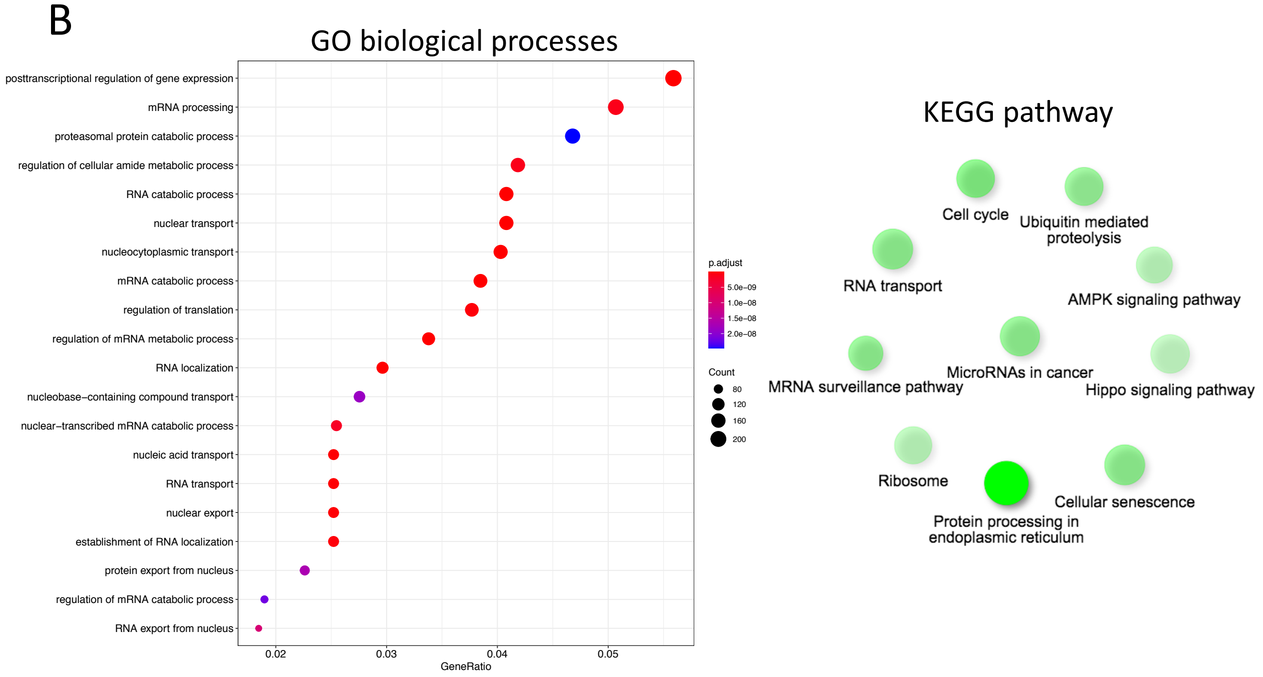


C

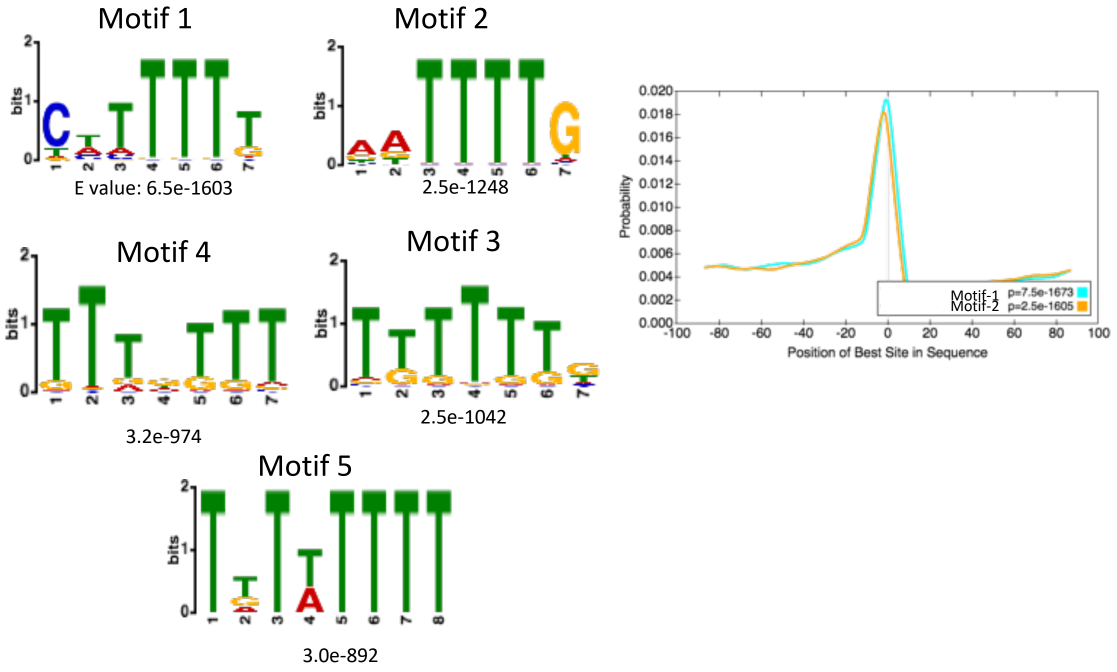

Extended Figure 4

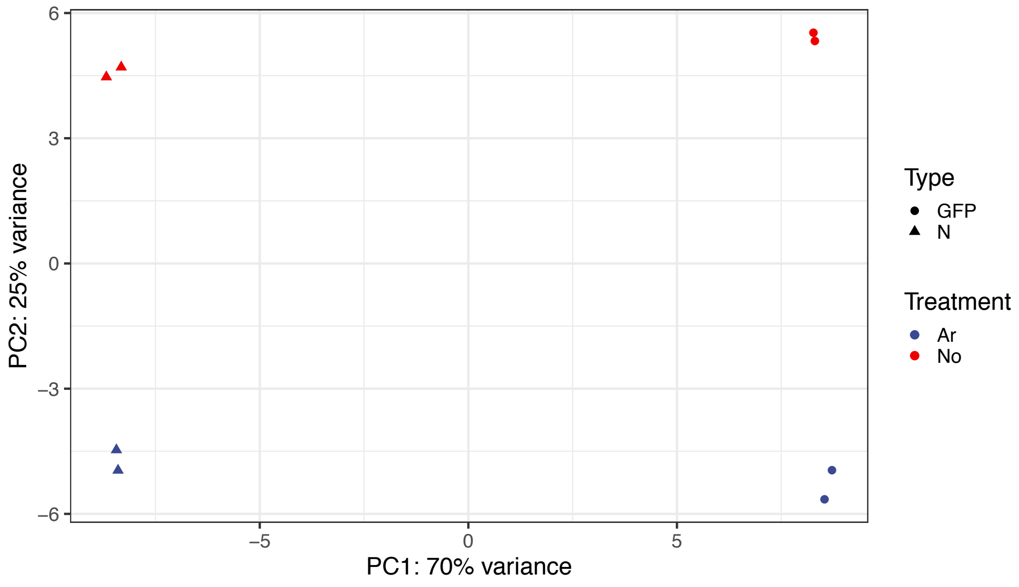

Extended Figure 5

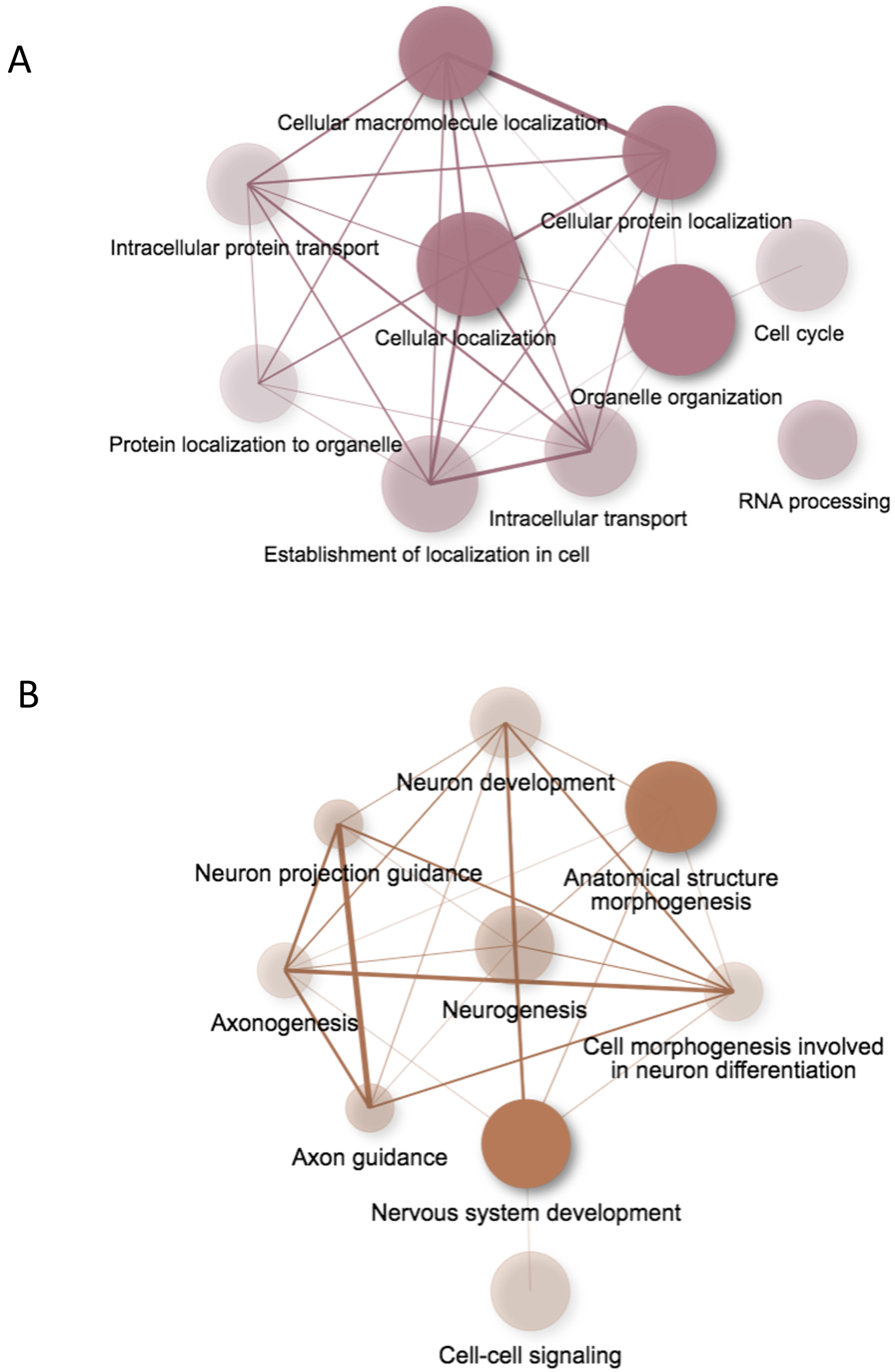

Extended Figure 6

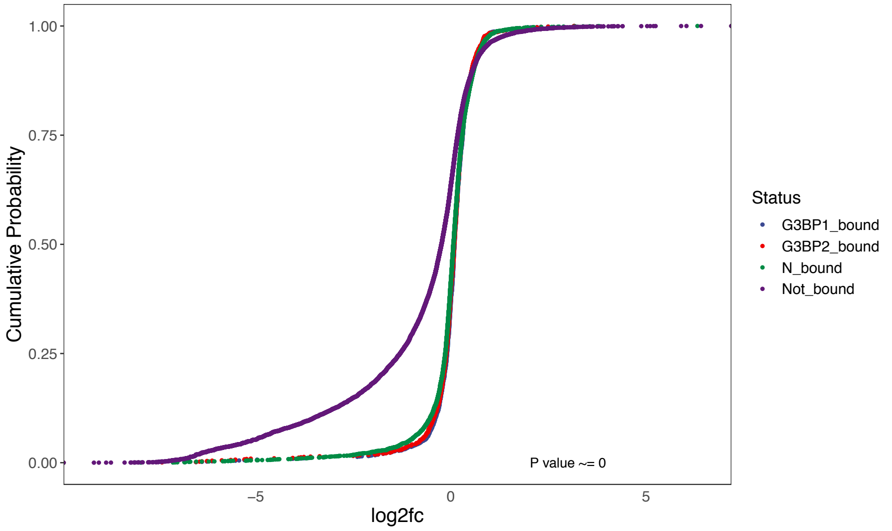
